# Acute effects of passive listening to Indian musical scale on blood pressure and heart rate variability among healthy young individuals – a randomized controlled trial

**DOI:** 10.1101/2020.05.03.073916

**Authors:** Kirthana Kunikullaya Ubrangala, Vijayadas, Radhika Kunnavil, Jaisri Goturu, Vadagenahalli S Prakash, Nandagudi Srinivasa Murthy

## Abstract

**Background:** Listening to music is entertaining but also has different health benefits. Music medicine involves passive listening to music, while music therapy involves active music making. Indian music is broadly classified into Hindustani and Carnatic music, each having their own system of musical scales (*ragas*). Scientific studies of Indian music as an intervention is meagre. Current study determines the effect of passive listening to one melodic scale of Indian music on cardiovascular electrophysiological parameters.

**Methods:** After informed consent, healthy individuals aged 18 – 30 years, of either gender were recruited and randomly divided into 2 groups (n=34 each). Group A was exposed to passive listening to the music intervention [Hindustani melodic scale elaboration (*Bhimpalas raga alaap*)], while group B received no intervention except for few natural sounds (played once in every 2 minutes). Blood pressure (BP, systolic – SBP; diastolic – DBP) and Electrocardiogram in lead II were recorded with each condition lasting for 10 minutes (pre, during, post). Heart rate variability (HRV) analysis was done. Data was analysed using SPSS 20.0 version and p<0.05 was considered significant.

**Results:** Passive listening to the musical scale employed had a unique effect. In group A, the SBP did not change during the intervention but increased insignificantly after the intervention was stopped (P=0.054). The DBP increased in both the groups during intervention and was significant among subjects in group A (P=0.009), with an increase of 1.676 mm Hg (P=0.012) from pre-during and 1.824 mm Hg (P=0.026) from pre-post intervention. On HRV analysis mean NN interval increased and HR reduced in both the groups, but was significant only in group B (P=0.041 and 0.025 respectively). In group A, most of HRV parameters reduced during music intervention, and tended to return towards baseline after intervention, but was statistically significant for Total Power (P=0.031) and Low Frequency (P=0.013) change; while in group B a consistent significant rise in parasympathetic indicators [SDNN, RMSSD, Total power and HF (ms^2^)] over 30 minutes was observed.

**Conclusion:** Unique cardiovascular effects were recorded on passive listening to a particular Indian music melodic scale, *raga Bhimpalas*, wherein, a mild arousal response, was observed. This could be due to attention being paid to the melodic scale as it was an unfamiliar tune or due to certain notes of this melodic scale, that particularly caused an arousal or excitation response. In contrast, the control group had only relaxation response. Exploring electrophysiological effects of different genres, melodic scales and its properties after familiarizing with the music may be illustrative.

## Introduction

Music is an aesthetic stimulus that has unique properties such as pitch, tempo, rhythm, scale, dynamic contrasts etc. The property, that when kept constant or varied, can specifically produce health benefits remains to be elucidated. Recent research recognized that specific acoustic factors within the musical signal induces a change in the processing of human cognitive and perceptual systems, to generate different bodily responses (1,2).

Indian music is broadly classified into Carnatic (South Indian) and Hindustani (North Indian) music, each with its own unique style. Music in India is said to have originated from Vedic chant tradition and traditional Persian music, as per ancient Indian music texts (3). *Ragas*, or musical or melodic scale, are permutations and combinations of various notes, in a specific order, in order to produce a melody. *Raga* may also be defined as a series of tones with specific melodic motifs that when improvised results in expression of certain emotions and creates a specified aesthetic experience (4,5).

In music theory, an interval is the difference in pitch between two sounds. An octave is the interval between one musical pitch and another pitch, that is double its frequency. The basic set of tones and relationships between them, that are used in *ragas* are derived are the 12-tone octave divisions / chromatic scale (6). Each interval is a tone defined by the ratio of its fundamental frequency to the tonic *(Sa). Swara* / note implies a note in the successive steps of the octave (7). With just 3 notes/ *swara* during Vedic times, the number increased to 5 and later 7 notes (*saptaswara* – represented as *Sa, Ri, Ga, Ma, Pa, Dha* and *Ni*, equivalent to *Do, Re, Mi, Fa, So, La, Ti* of western music), which is now considered ideal to produce a melodic scale / *raga* (6,8). Each melodic scale is organized as *Aarohana* (ascending sequence of notes) and *Avarohana* (descending sequence), is further improvised, within the framework of the scale, in vocal or instrumental performances, presenting the various aspects of the scale (e.g., sustainance of notes, elaboration, timing, ending notes, repeated notes etc.). The ‘major’ intervals are the *shuddh swaras* or the natural notes namely, second, third, sixth, and seventh while the ‘minor’ intervals are the *komal swaras* (flat) positions of the same tones. Indian music improvisation has unique set of rules that is pre-determined but yet creative, and *alaap (vistar), jor, swarakalpana, taan, tanam, neraval* and so on, all form different parts of this improvisation. *Alaap* is quasi-creative improvisation, seen in both Carnatic and Hindustani music, where, note by note is elaborated, presenting the prominent phrases of the scale, usually beginning in a slow tempo, with progress to medium and faster tempos, but not bound by any rhythmic cycle (9–11). It is beyond the scope of this article to describe all types of improvisations.

Music not only has benefits that are psychological but physiological as well. It can regulate stress mechanism, sleep wake cycle, improve cognitive skills, beneficially effect the blood pressure (BP), heart rate (HR), respiration rate (RR), body temperature, and biochemical parameters as well as sensitivity to pain (12–17). Music literature as well as past studies have shown that specific *ragas* elicit distinct emotions / *rasas* (18–22). As a result of increased interest and research in this field, musical auditory stimulation is now proposed as a non-pharmacological intervention or as a complementary therapy (23–25).

One of the initial works exploring the cardiovascular effects of music, was in 1918, by Hyde *et al*, who found a decrease in systolic blood pressure (SBP) and diastolic blood pressure (DBP) when minor tones were used, whereas, the stirring notes of Toreador’s song increased the SBP and HR (26). Cardiorespiratory parameters were modified on repeated rhythmic recitation of a prayer, poetry or yoga mantra (27,28). Listening to sedative music (slow tempo, legato phrasing, and minimal dynamic contrasts) was shown to reduce HR and BP. BP was shown to be proportional to the crescendo present in music, whereas music with uniform emphasis reduced the BP (16). A study has shown that music was as effective as benzodiazepines in reducing BP (29).

Several studies reported that under various conditions music decreases sympathetic nervous system (SNS) and increases parasympathetic nervous system (PNS) activity as measured by HR and heart rate variability (HRV), indicating physiological relaxation (30–34). However, no difference in HR or HRV was observed by a few investigators (35,36), an increase was reported by some (32,37). A few works showed that music decreased Low frequency / High frequency (LF/HF) ratio (34), while a few others showed an increase in LF/HF (32,37). When music was intervened with randomly inserted 2 minute pauses it was observed that passive listening to music increases BP, HR and LF/HF in proportion to tempo and perhaps to the complexity of the rhythm. It was found that silence (pauses), that followed the music, induced more parasympathetic stimulation. One Indian track (*raga Maru Behag* played on Sitar) used by the authors, which had a tempo of 55 beats/min, however, induced a significant large fall in HR (32). People with different tastes in music respond differently, and those who are not involved in the music have no response (38). One study, where use of *Rabindra sangeet* improved HRV, the authors hypothesized that the effect differed from person to person (39). A recent systematic review also concluded that music does have positive effects on autonomic nervous system (12).

We thus observe that there is varying literature available on the effects of music on the cardiovascular parameters and the mechanisms behind it. However, there are very few studies that have used Indian music scientifically as an intervention for health benefits. In India, music is predominantly used as entertainment. Despite ample vedic literature available on the beneficial effects of melodic scales / *ragas* on human mind and body, scientific evidence for the same is extremely meagre. In our previous work we showed that BP reduced significantly after listening to Indian music among prehypertensives. All subjects were given a musical piece composed on *raga ‘bhimpalas’* (*raga* that is said to normalize BP(40)) to be heard daily, for 15 minutes a day, for at least 5 days a week, for 3 months. Here, 24 hour ambulatory BP and HRV were recorded once on recruitment and followed up after 3 months. On retrospection into our methodology, the acute effect of passive listening to the *raga* on BP or HRV was not explored, nor the effect of other *ragas* listed in *Gandharva veda* (that could normalize the BP) were scientifically evaluated, for their electrophysiological effects (15,41).

The hypothesis of the present study was that, an Indian musical scale, the *Hindustani raga, Bhimpalas*, would reduce BP and increase parasympathetic activity analysed through HRV, during a passive listening task, that would return to baseline after intervention, among young healthy individuals.

### Methodology

A double blinded, prospective, randomized controlled trial was conducted with an experimental study design, with a total sample of 68, randomized into 2 groups (n=34 subjects in each group). Group A was exposed to passive listening to the music intervention [Hindustani melodic scale elaboration (*alaap*)], while group B received no intervention except for few natural sounds (played for 10 seconds once in every 2 minutes to avoid sleeping). The study protocol was approved by the institutional scientific committee on human research and ethical review board.

### Basis for sample size

The sample size was calculated (using nMaster 2.0 sample size software, Department of Biostatistics, CMC, Vellore) based on a study conducted by Okada *et al* (34) it was found that RMSSD (root mean square standard deviation of NN intervals on ECG) was 17.4 (7.2) ms and 24.1 (15.5) ms before and after music therapy. With an effect size of 0.59 and power of 90% and confidence interval of 95%, the minimum sample size required for the present study was estimated to be 32, in each group.

### Recruitment of subjects for the study

Ramaiah group of institutions comprise of people from medical, dental, pharmacy, physiotherapy, engineering etc. backgrounds. Healthy subjects aged 18 – 30 years were invited to participate in the study via advertisements in notice boards of various institutions, social media posts and posters. Participants who responded to the call were sent an online questionnaire via google forms, as explained further. About 100-120 responded to the call, of which 80 completed the google form. Inclusion criteria were healthy subjects, aged 18-30 years, of either gender, non-smokers and alcoholics. Exclusion criteria was any medical disorder (cardiovascular, renal, respiratory, endocrine, hearing problem, psychiatric disorders, stroke, epilepsy), pregnancy, body mass index (BMI)>30 kg/m^2^; intake of drugs which are known to affect the BP or autonomic status of the individual, other impairments that would prevent the subject from performing few experimental procedures. The healthy cardiovascular system of the volunteers was defined by measuring BP, that confirmed their non-hypertensive state and by measuring baseline HR that confirmed their non-tachycardiac state.

### Baseline demographic data collection

A pre-tested, pre-designed web-based questionnaire (Google forms) was implemented, so that it is convenient for the subjects enrolled to enter their data easily. This questionnaire contained details such as subject’s name, gender, socio-demographic details, education background, drug history, present or past history of non-communicable diseases if any and family history of non-communicable disorders, smoking and alcohol history. A few questions inquiring the subjects’ preference to any type of music, previous experience with music (instrumental or vocal) was also included. Following the collection of data online, the subjects were invited, for further data collection, to the lab.

Out of 80 subjects who answered the online questionnaire, 75 subjects reported to the lab. The subjects were interviewed and all the information was collected after establishing rapport with them. After overnight fasting, they were asked to take a light breakfast and abstain from exhaustive exercise, for the past 24 hrs. They were asked to abstain from tea, coffee about 2 hours prior to the recording. A general health check-up was done for all subjects. The BMI was calculated and BP in sitting position was measured twice after five minutes’ rest (Sphygmomanometer) in between, and was noted (42). Only normotensives were included as per inclusion criteria. Recruited subjects (n=68) were explained about the study protocol, and co-operation expected from them and informed consent was obtained to participate in the study. They were informed about their rights to withdraw their participation from the study.

### Randomization

All subjects were randomized into 2 groups using simple randomization technique. The random numbers were computer generated using MS Excel (2 sets of 34 each). The random number indicating intervention or control was kept in an opaque and sealed envelope and the serial number of the subjects were written on the top of the envelope. The envelope was opened by the research assistant after the baseline assessment of each participant had been completed and assigned the participants randomly to both the arms, into intervention and control categories. All the investigators who did the outcome assessments were blinded to the interventions.

### Baseline (Pre) and Post intervention readings

All the recordings were carried out between 08:00 am and 10.00 am in an isolated examination room at a stable temperature between 20 and 22°C, in a noise free atmosphere. It was ensured no one entered the lab once recordings began. The subjects were asked to relax in a bed for about 10 minutes prior to the tests, with their eyes closed. They were asked to remain as still as possible to exclude movement induced artefacts, and also refrain from talking, falling asleep and intentionally altering their respiration during the recording. We also instructed the subjects to breathe at a rate of 6 breaths per minute, throughout the procedure that was monitored on the computer software. Subjects were carefully monitored to ensure there were no significant respiratory or postural changes during the session.

During the first ten minutes, BP cuff was tied to the left arm of the subjects and one reading was taken for the subject to know the feel of automatic cuff inflation and deflation. Recording of BP, was done using digital BP monitor as a normal sphygmomanometer recording would not only disturb the subject during the intervention but also delay the recordings and eliminate the effect the intervention. A standardized digital BP monitor was used (Omron HEM-7130L, Europe), the reliability of which has been established (43). Electro-cardiogram (ECG) was recorded in Lead II (sample rate of 1000 Hz) for ten minutes, as it is twice the minimum window required for HRV analysis. The recording of the data began in the Power lab 15 T Lab chart hardware & software (AD instruments).

After all the attachments, within the first 10 minutes, baseline ECG recording commenced. At the end of 10 minutes, baseline digital measurement of BP (systolic, diastolic BP and pulse rate) was done and recorded as pre-intervention readings. This event was marked in ECG and the recordings continued. After this music intervention began, and the event was marked. At the end of 10 minutes of music, without disturbing the subject the BP was recorded. ECG monitoring was continued for another 10 minutes and at the end, the event was marked. Post intervention BP was recorded and the subjects were made to feel comfortable and were relieved.

### Intervention

The 2 mp3 recordings were coded as A and B by a person uninvolved in the present study. We instructed the subjects to listen to this with eyes closed, mind relaxed, for the duration it was played. The subjects listened to the music through headphones [studies have previously used headphones, which is considered ideal as per the review (44)], connected to a laptop, at uniform volume (50%).

#### Music intervention

For music intervention, the previously standardized melodic scale *Bhimpalas* was used for the present study (15,41). It contained instrumental (*Bansuri*) music recorded by an eminent flautist, playing the respective *alaap* in the *raga* (musical scale). The melodic scale / *raga* was played for 10 mins duration. The subjects in Group A listened to music.

*Bhimpalas raga*, belongs to the *Kapi thaat*, is a soft, poignant and passionate *raga* that evokes a feeling of love and yearning. It is generally classified as a ‘late-afternoon’ *raga*. In Carnatic music (South Indian classical music) the *raga* ‘*Abheri*’ is the closest counterpart of this Hindustani *raga* (45). The scale of this *raga* is as follows: *Aroha?a:* S G_2_ M_1_ P N_2_ S. *Avaroha?a:* S N_2_ D_2_ P M_1_ G_2_ R_2_ S (*shadja, shuddha Rishabh, komal Gandhar, suddha Madhyam, pancham, shuddha Dhaivath, komal Nishadh*). Equivalent notes on western scale are B♭ C E♭ F G B♭ C as ascent and C B♭ A G F E♭ D C as descent on western scale (46). Thus the scale is made up of two flat keys and no sharps.

#### Intervention to the control group

The control group (Group B) did not receive any intervention, but since the complete recording lasted for 30 – 40 minutes’ duration, it was possible for the subjects to feel sleepy or fall asleep. Sleep would cause its own electrophysiological effects, which would alter the objective of the present study. Further, silence during the middle 10 minutes would not be an ideal to compare, when the other group received music. For these 2 reasons, natural sounds (birds chirping and flowing river) was played for 10 seconds duration once in every 2 minutes in the mid-10 minutes (intervention phase).

### BP Analysis

The readings given by the BP monitor were recorded as SBP (in mm Hg), DBP (in mm Hg) and HR (in beats per minute). The readings were taken before (end of 10 minutes of relaxation – pre-intervention), during (end of 10 minutes of intervention) and after (end of 10 minutes after the intervention was stopped) intervention.

### HRV Analysis

Of the whole recording the first 1-2 minutes of each segment of data were excluded in case of any transition or adjustment effect. Only series with more than 95% of sinus beats was used for analysis. Time domain paramters analysed using fast Fourier transformation (FFT size: 1024) were SDNN—the standard deviation of NN intervals, RMSSD—Root square of the mean squared difference of successive NNs, NN50—number of pairs of successive NNs that differ by more than 50 ms, pNN50—proportion of NN50 divided by total number of NNs, spectral components such as Very Low Frequency (VLF), Low Frequency (LF) and High Frequency (HF) components in absolute values of power (ms^2^) and in normalized units (nu), and LF/HF.

A region in the channel that contained data without much variation and ectopics was selected and analysed. A threshold value was set to detect the beats (R waves – R component of QRS complex) and it was increased to avoid detection of unwanted peaks or decreased to detect genuine beats that would have been missed. Beats which fall outside of the timing of a normal sinus rhythm were considered ectopics. Ectopics were excluded as they do not represent ANS activity and are not believed to contribute to HRV. Inclusion of ectopics during analysis results in falsely higher representation of HF component of HRV (47,48). Poincare plot, where RR interval is plotted against the preceding RR interval, in a scatter plot analysis, has been widely used as a quantitative visual tool for HRV analysis (49–51). After referring the Poincare plot (for the best possible ellipse), RR interval Tachogram, a plot of successive RR interval values against the interval number, and the spectrum for any ectopics and detection of R waves, a report was generated and results were entered onto an excel sheet and tabulated. Sources of error were minimized by only having one of the investigator perform the recording of ECG and analysis of HRV of the subjects. The paramters analysed were pre, during and post intervention mean NN interval, HR (Average of 10 minutes), SDNN, RMSSD, NN50, pNN50, VLF, LF, HF (ms^2^), LF nu, HF nu and LF/HF.

### Statistical analysis

Data was analysed using SPSS software version 18.0 (SPSS Inc. Released in 2009. PASW Statistics for Windows, Version 18.0. Chicago: SPSS Inc.). The continuous variables were analysed using descriptive statistics using mean and SD. The categorical variables were analysed using frequency and percentage. The normalcy of the data was checked by applying the Kolmogorov-Smirnov Test. All the variables namely BP, HR, and HRV were found to follow the normal distribution.

Baseline comparisons between the groups were carried out using students’ t-test for continuous variables and chi-square test for categorical variables. All HRV parameters were compared between groups and at pre, during and post-intervention using repeated measures of ANOVA (RM ANOVA) test. Bonferroni multiple comparisons test was used to compare the pairwise differences. All the baseline parameters were comparable between the groups. Only baseline DBP showed a significant difference between the groups and hence multivariate regression analysis and stratified analysis were carried out to adjust for the various covariates. Two-tailed P value <0.05 was considered for statistical significance.

## Results

A total of 68 subjects were enrolled into the study, with each group consisting of 34 subjects. The two groups were comparable based on mean age, age distribution and gender. There were more subjects in the age group of 19-21 years in both the groups. Groups were comparable with respect to BMI with statistically no significant difference (P=0.307).

About 30% of individuals out of 68 recruited were musically trained, with about 29.4% in group A and 32.4% in group B (P=0.793). Subjects were predominantly trained in Indian music (91.7%). Most of the subjects preferred listening to old *Hindi* movie songs.

Baseline comparison of the parameters were carried out between the 2 groups, which revealed that SBP (P=0.501), HR (P=0.8) and all the parameters of HRV were comparable. However, DBP showed statistically significant difference between the groups (P=0.003).

### Before intervention

All sociodemographic and baseline parameters were comparable between the two groups, except DBP (P=0.010), which was higher in the control group prior to intervention. However, on regression analysis of all the variables (age, gender, education, diet, marital status, involvement in physical activity, mind body relaxation techniques, family history of non-communicable disorders, training in music, preference to music along with differences in BP based on conditions), none of the parameters seemed to affect the change in DBP that was observed.

### BP During and after intervention

The key findings in BP at 20^th^ minute (during music) in group A were - SBP was similar to pre-intervention levels during intervention and increased after intervention (P=0.054). When the differences amongst the groups were tested, statistically no significant difference was found (P = 0.259). The DBP increased significantly (P=0.009), by 1.676 mm Hg (P=0.012) during intervention and by 1.824 mm Hg; (P=0.026) after intervention in comparison to pre-intervention levels. The HR insignificantly reduced during and increased after intervention (P=0.937). In control group, SBP reduced during intervention and after intervention (P=0.22), DBP increased during intervention and later reduced (P=0.403). The HR continued to reduce throughout the 30 minutes’ duration (P=0.74) [Table 3; Figure 2]. Note that the HR measured using BP apparatus was recorded one time along with BP and was not an average of continuous monitoring before, during or after intervention.

**Table 1:** Scale of Raga Bhimpalas, the names of the notes in Hindustani music and Western scale, with their equivalent frequencies, just intonations and 12-TET

| Svara /<br>Note | Hindustani name | Staff<br>note | Western scale<br>Interval name | Frequency | Just<br>intonation<br>(Cents) | 12-TET<br>(Cent) |
| --- | --- | --- | --- | --- | --- | --- |
| Sa | Shadja | C | Perfect unison | 1 | 0 | 0 |
| re | Shuddha rishab | D | Major second | 10/9 | 183 | 200 |
| ga | Komal gandhar | E<sub>b</sub> | Minor third | 6/5 | 316 | 300 |
| Ma | Shuddha madhyam | F | Perfect fourth | 4/3 | 498 | 500 |
| Pa | Pancham | G | Perfect fifth | 3/2 | 702 | 700 |
| Dha | Shuddha daivat | A | Major sixth | 5/3 | 884 | 900 |
| ni | Komal nishad | B<sub>b</sub> | Minor seventh | 9/5 | 1018 | 1000 |

Note: Interval names, abbreviations, frequency ratios, and sizes in cents for just intonation (JI) as well as 12-tone equal temperament (12-TET) tunings are shown. The 12 intervals of the Western chromatic scale, comparably presented (More about JI and 12-TET in supplementary file).

**Table 2:** Distribution of subjects under music intervention group (Group A) and control group (Group B).

| Variables | Group | Group A<br>n (%)<br>N=34 | Group B<br>n (%)<br>N=34 | Total<br>n (%)<br>N=68 | P -Value |
| --- | --- | --- | --- | --- | --- |
| Age (Years) | <=18 | 7 (20.6) | 4 (11.8) | 11 (16.2) | 0.651 |
|  | 19-21 | 19 (55.9) | 19 (55.9) | 38 (55.9) |  |
|  | 22-24 | 4 (11.8) | 4 (11.8) | 8 (11.8) |  |
|  | >=25 | 4 (11.8) | 7 (20.6) | 12 (16.2) |  |
| Age (years)<br>Mean, SD |  | 20.3, 2.60 | 21.0, 2.71 | 20.8, 2.80 | 0.278 |
| Gender | Female | 16 (47.1) | 9 (26.5) | 25 (36.8) | 0.078 |
|  | Male | 18 (52.9) | 25 (73.5) | 43 (63.2) |  |
| Mean BMI<br>(kg/m<sup>2</sup>)<br>Mean, SD |  | 23.4, 4.67 | 22.3, 4.02 | 23.0, 4.45 | 0.307 |
| Training in<br>Music | No | 24 (70.6) | 23 (67.6) | 47 (69.1) | 0.793 |
|  | Yes | 10 (29.4) | 11 (32.4) | 21 (30.9) |  |
| Genre of<br>music | Indian | 9 (90.0) | 13 (92.9) | 22 (91.7) | 0.803 |
|  | Western | 1 (10.0) | 1 (7.1) | 2 (8.3) |  |

**Table 3:** Comparison of absolute values and logarithmic levels of BP (in mm Hg), HR and HRV in between 2 groups, pre, during and post intervention

|  | Group A (N=34) |  |  |  | Group B (N=34) |  |  |  | P (between groups)** |
| --- | --- | --- | --- | --- | --- | --- | --- | --- | --- |
|  | Pre | During | Post | P within group | Pre | During | Post | P within group |  |
| **SBP (mm Hg)<sup>a#</sup>** | 107.6, 12.79 | 107.4, 13.96 | 109.4, 13.20 | 0.054 | 105.7, 9.98 | 105.4, 10.26 | 103.8, 11.43 | 0.22 | 0.259 |
| **DBP (mm Hg)<sup>a#</sup>** | 67.2, 6.10 | 68.9, 7.37 | 69.1, 6.51 | **0.009\*** | 71.8, 6.09 | 73.0, 6.82 | 72.4, 7.83 | 0.403 | **0.010\*** |
| **HR (bpm)<sup>a#</sup>** | 71.91, 8.5 | 71.68, 9.26 | 72.03, 7.01 | 0.937 | 72.59, 12.94 | 72.18, 8.81 | 71.62, 9.64 | 0.74 | 0.903 |
| **Mean NN (ms)** | 2.9, 0.04 | 2.9, 0.05 | 2.9, 0.04 | 0.091 | 2.9, 0.07 | 2.9, 0.07 | 2.9, 0.06 | **0.041\*** | 0.541 |
| **HR (bpm)<sup>a^</sup>** | 72.9, 7.09 | 72.1, 7.26 | 71.7, 7.18 | 0.061 | 72.8, 11.29 | 71.1, 10.49 | 71.0, 10.59 | **0.025\*** | 0.778 |
| **SDNN** | 1.8, 0.17 | 1.7, 0.17 | 1.7, 0.21 | 0.405 | 1.8, 0.18 | 1.8, 0.17 | 1.8, 0.18 | **0.004\*** | 0.231 |
| **RMSSD** | 1.7, 0.26 | 1.7, 0.26 | 1.7, 0.26 | 0.337 | 1.7, 0.28 | 1.8, 0.24 | 1.8, 0.25 | **0.040\*** | 0.384 |
| **NN50** | 2.2, 0.57 | 2.2, 0.48 | 2.2, 0.46 | 0.802 | 2.2, 0.58 | 2.2, 0.50 | 2.2, 0.51 | 0.278 | 0.914 |
| **pNN50** | 1.3, 0.52 | 1.3, 0.48 | 1.3, 0.49 | 0.864 | 1.3, 0.62 | 1.3, 0.54 | 1.4, 0.37 | 0.128 | 0.412 |
| **TP (ms<sup>2</sup>)** | 3.5, 0.35 | 3.4, 0.34 | 3.5, 0.36 | **0.031\*** | 3.5, 0.38 | 3.5, 0.35 | 3.6, 0.37 | **0.009\*** | 0.348 |
| **VLF (ms<sup>2</sup>)** | 3.0, 0.33 | 2.9, 0.31 | 2.9, 0.40 | 0.173 | 2.9, 0.38 | 3.0, 0.29 | 3.0, 0.34 | 0.113 | 0.336 |
| **LF (ms<sup>2</sup>)** | 2.8, 0.37 | 2.8, 0.38 | 2.9, 0.37 | **0.013\*** | 2.9, 0.39 | 3.0, 0.38 | 3.0, 0.35 | **0.025\*** | 0.084 |
| **LF (nu)** | 1.6, 0.20 | 1.5, 0.25 | 1.6, 0.22 | 0.269 | 1.6, 0.17 | 1.6, 0.19 | 1.6, 0.19 | 0.573 | 0.275 |
| **HF (ms<sup>2</sup>)** | 3.0, 0.45 | 3.0, 0.44 | 3.0, 0.46 | 0.555 | 3.0, 0.50 | 3.1, 0.43 | 3.1, 0.46 | **0.023\*** | 0.527 |
| **HF (nu)** | 1.7, 0.16 | 1.7, 0.17 | 1.7, 0.16 | 0.233 | 1.7, 0.19 | 1.7, 0.16 | 1.7, 0.15 | 0.387 | 0.79 |
| **LF/HF** | 0.9, 0.32 | 0.9, 0.36 | 1.0, 0.33 | 0.206 | 0.9, 0.31 | 0.9, 0.36 | 1.0, 0.34 | 0.682 | 0.751 |

Note:
- N is the number of subjects in each group. - <sup>a</sup>absolute level #one-time measurement; <sup>^</sup>Average of continuous monitoring over 10 minutes - BP is given in (mm Hg) and HRV all parameters have been log converted, except HR, which is in beats per minute (bpm) - For all HRV parameters, the null hypothesis (H<sub>0</sub>) considered was that mean values is the same at all the time points (pre, during and post). The alternative hypothesis is that mean value is significantly different at one or more time points. - For absolute values of all HRV parameters refer supplementary file - All the values are in mean, standard deviation (SD) – univariate ANOVA. - P value < 0.05 was considered significant – Levene's test of equality - \*\*P calculated using RM-ANOVA.

**Figure 1:**
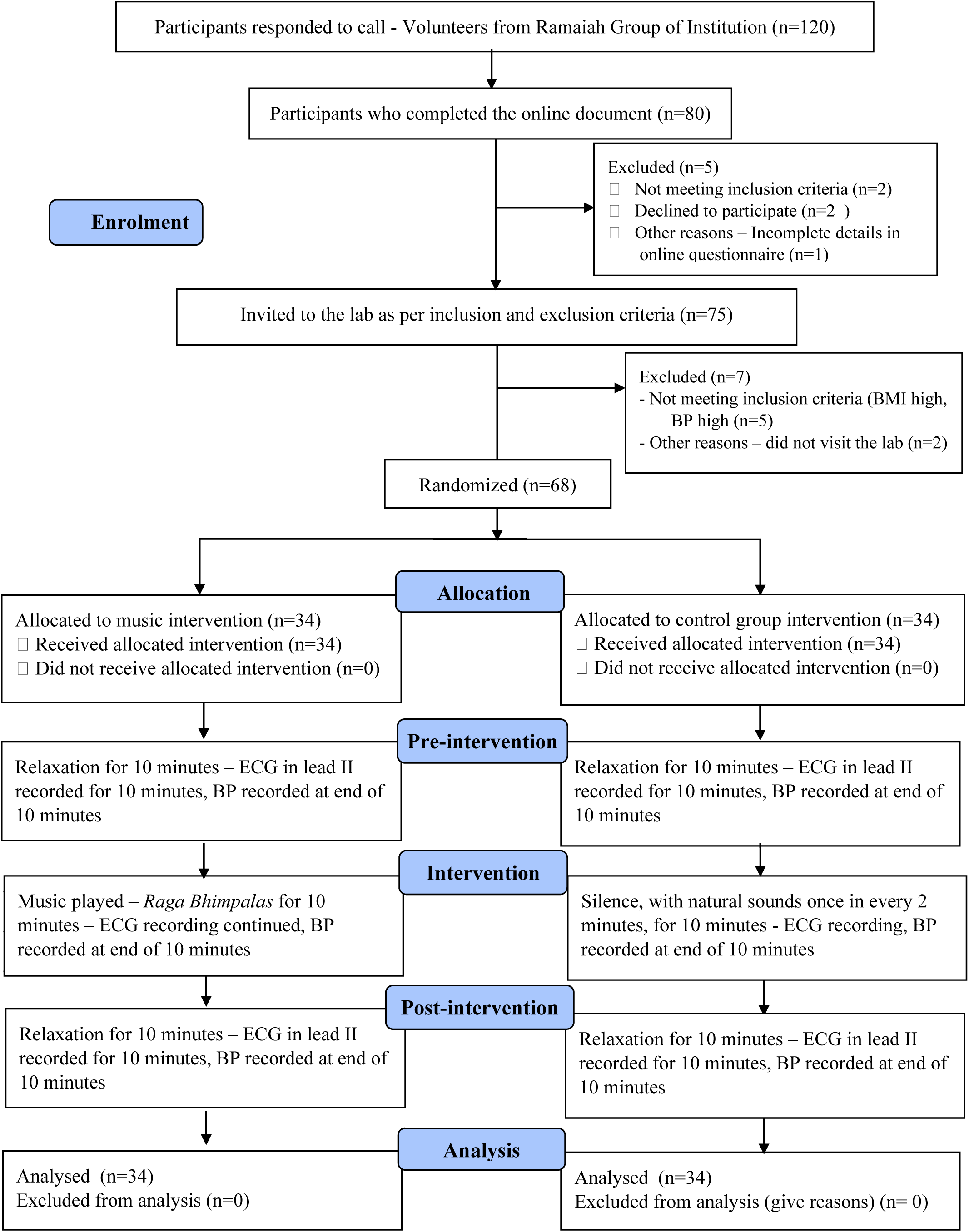
Consort diagram of participant recruitment, random allocation, data collection and follow up.

**Figure 2:**
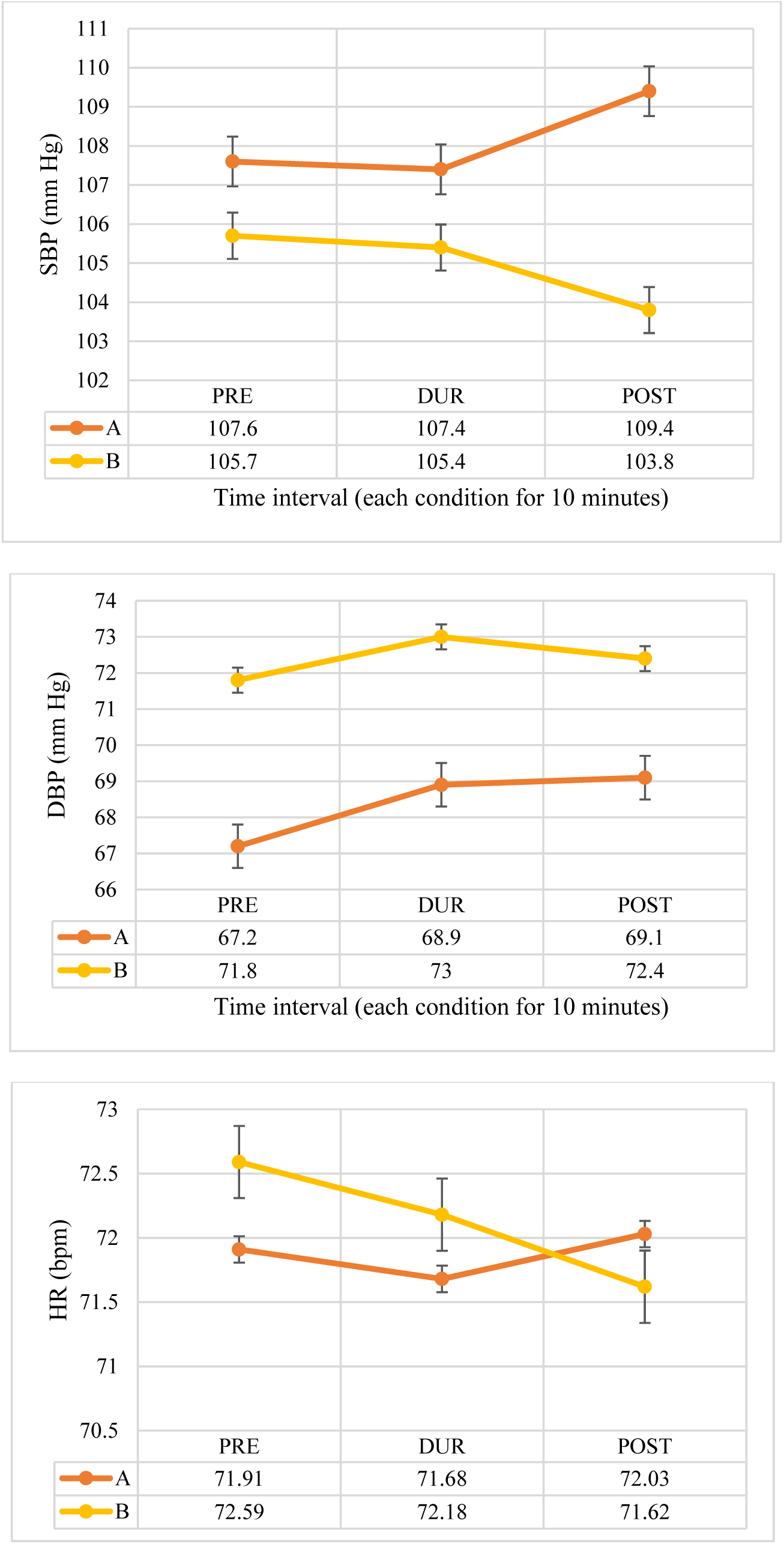
Comparison of SBP, DBP (in mmof Hg) and HR (in bpm) between 2 groups (Pre, Dur and Post intervention)

**Figure 3:**
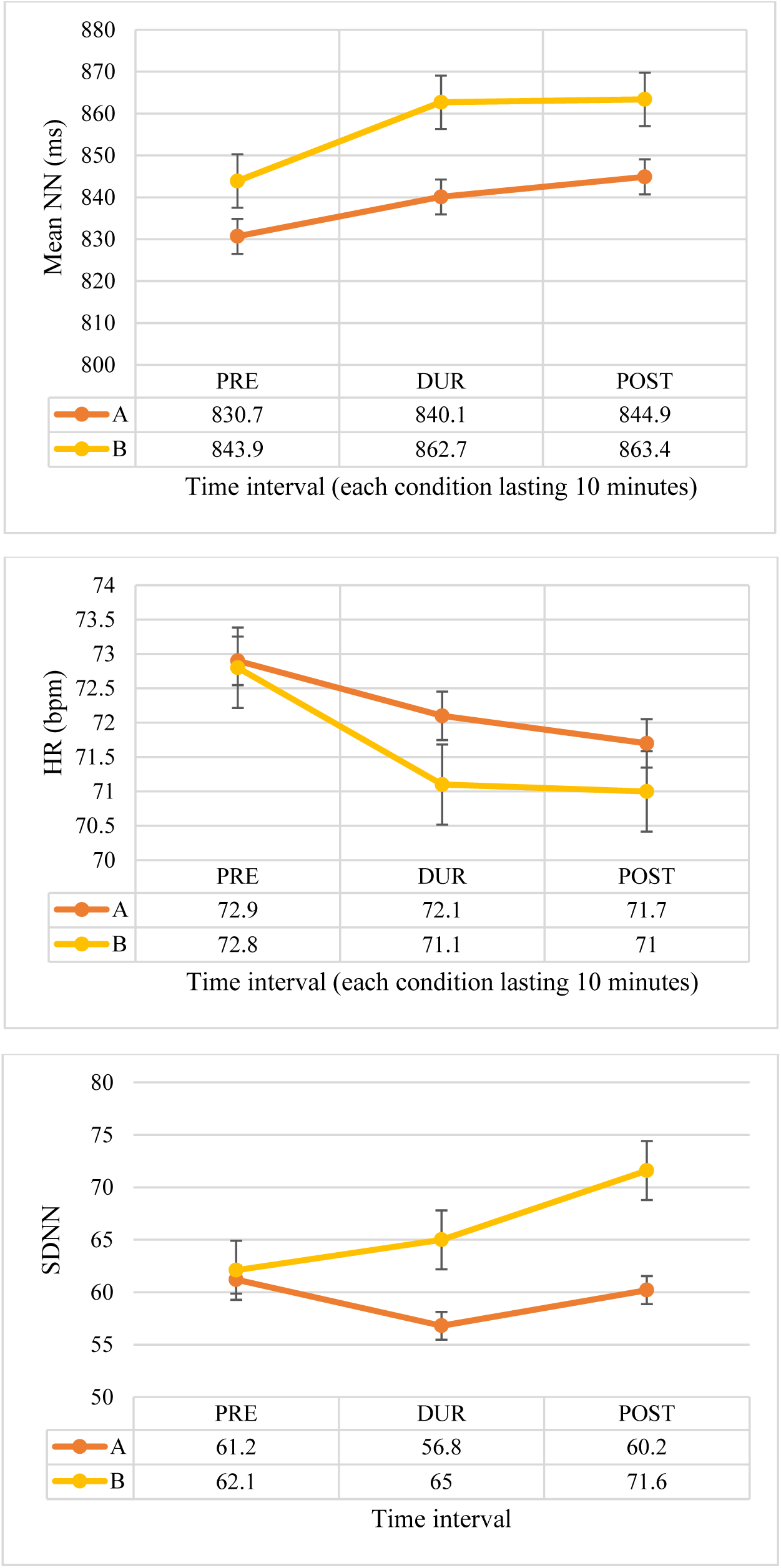
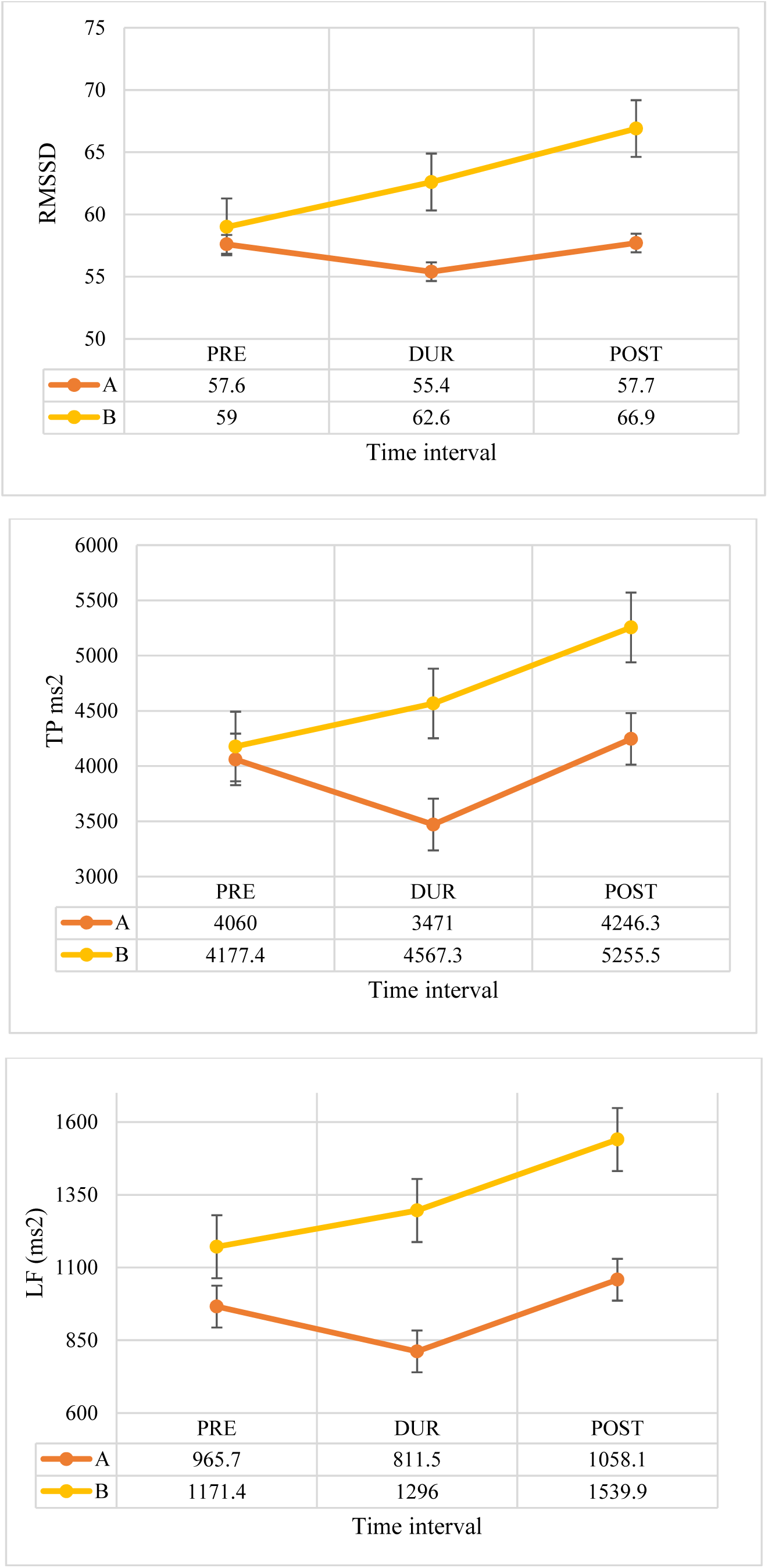
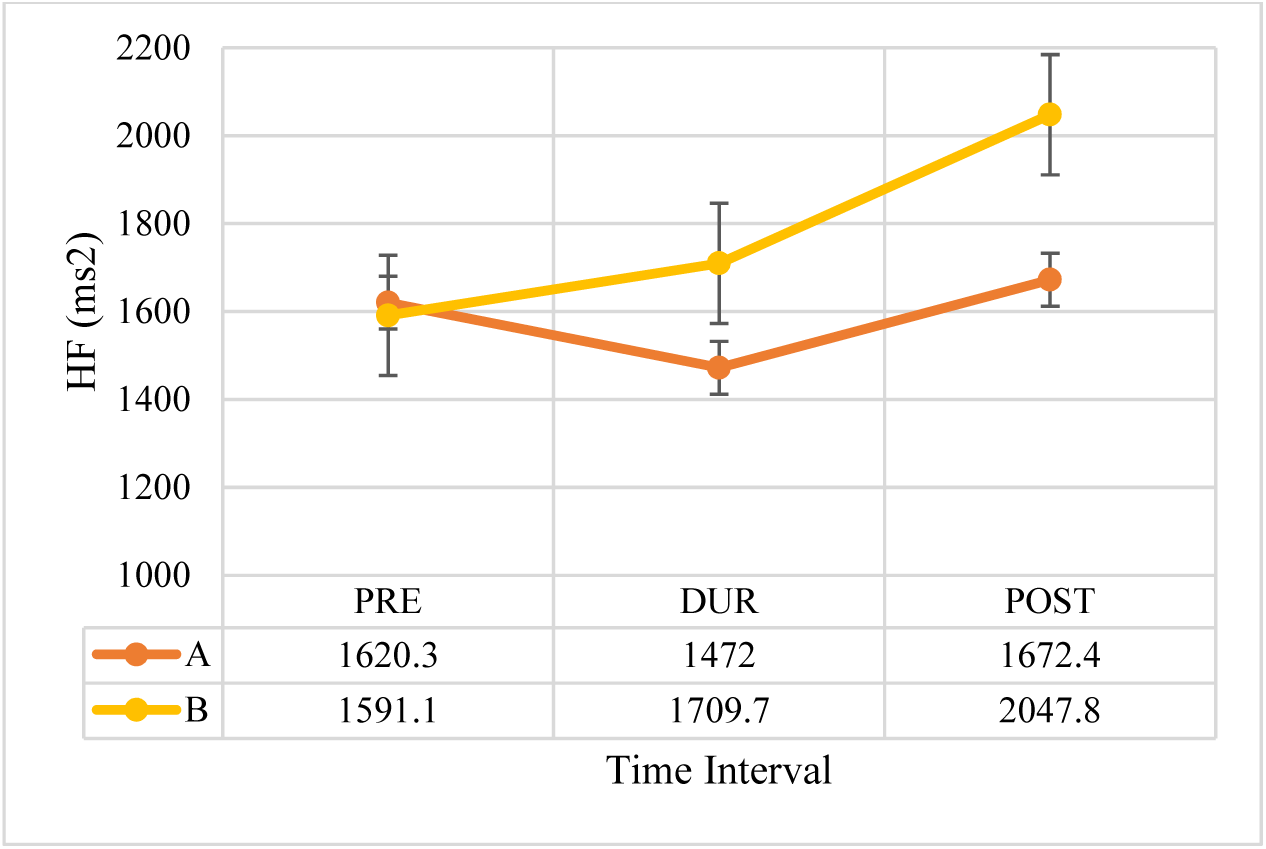
Comparison of Absolute values of Mean NN Interval (ms), HR (Lead II ECG, Averaged), SDNN, RMSSD, TP(ms2), LF(ms2) and HF (ms2) between 2 groups (Pre, Dur and Post intervention). Note that the absolute values have been given in data table below each graph. For all absolute values, refer supplementary file.

### HRV During and after intervention

Note that the changes in HRV were the average of 10.1^th^ to 20^th^ minute of ECG analysis. For HRV analysis, all parameters except HR were log transformed due to skewness in data obtained (absolute levels of all HRV parameters and pairwise comparison has been shown in Supplementary file). ANOVA with repeated measures^1^ was used with sphericity assumption. Further, two way repeated measures of ANOVA analysis was done to inspect the interaction between intervention group and time. The results are as follows.

In group A, during intervention, the various parasympathetic parameters of HRV [SDNN, RMSSD, TP, VLF, LF, HF (ms^2^)] reduced along with HR, but was statistically not significant. The mean NN interval increased, but not significantly (P=0.091). After intervention mean NN interval continued to increase, HR reduced, while SDNN, RMSSD, TP (ms^2^), HF (ms^2^) and LF (ms^2^) increased towards pre-intervention levels. The change was significant for TP (ms^2^) (Global HRV) (P=0.031) and LF (ms^2^) (P=0.013). On pairwise comparison LF ms2, the change was maximal after intervention compared to during music (P=0.005). Though NN50 and pNN50 reduced in group A, after log transformation the change was statistically not significant.

In control group, during and after intervention, sustained and significant increase (P≤0.05) in mean NN interval (P=0.041), SDNN (P=0.004), RMSSD (P=0.040), TP (ms^2^) (P=0.009), LF (ms^2^) (P=0.025) and HF (ms2) (P=0.023) was observed along with reduced HR reduced (P=0.025). On pairwise comparison, maximum change in Mean NN interval was observed from pre-intervention to during intervention interval (P=0.005). The reduction in HR was maximum during intervention compared to baseline levels (P=0.002). The unit drop in HR was very less (hardly 1 beat per minute). SDNN change was significant after intervention compared to baseline (P=0.014) and during (P=0.024) levels. Total power change was significant after intervention compared to baseline levels (P=0.026). The change in LF (ms^2^) was maximal after intervention, compared to during (P=0.049) intervention. These changes observed in HRV parameters in its absolute power were statistically not significant after normalized unit (nu) conversion.

## Discussion

In this study, to the best of our knowledge, for the first time, an Indian musical scale (*Hindustani raga)* has been evaluated scientifically for its effect on electrophysiological parameters such as BP and HRV in young, clinically normal, normotensive individuals.

Vedic literature (*sama veda*) and *raga chikitsa* literature specifies about *7 ragas* to be useful in BP control (*Ahir bhairav, Kausi Kanada, Bhimpalas, Todi, Puriya, Hindol and Bhupali)*. In our previous study we observed that *Bhimpalas raga* could effectively control DBP among prehypertensives, after 3 months of music intervention (41). Though we found noticeable reduction using single *raga Bhimpalas* music intervention, along with standard management protocols, there was not enough scientific evidence regarding the acute effect of the *raga* on normal healthy individuals, be it on BP or on other electrophysiological parameters. Therefore, in the current study, previously standardized melodic scale, *raga Bhimpalas*, was scientifically evaluated, for its acute effects.

Those in the intervention arm (group A), passively listened to *bansuri* (Indian flute) recording containing only *alaap* (a type of improvisation in Indian music) in *raga Bhimpalas*, against the drone instrument in the background, without percussion instruments, through headphones connected to a laptop for 10 min. The BP and HRV was digitally monitored thrice [pre, during and post – each condition lasting for 10 min]. Those in control arm (group B) relaxed for 30 minutes, when the physiological parameters were recorded. To make the control group matched for the intervention and to avoid sleeping (and its effects), control arm received acoustic stimuli with natural sounds lasting for 10 seconds, played once in every 2 minutes, during intervention condition (mid 10 minutes).

### Main findings

#### Before intervention

All sociodemographic and baseline parameters were comparable between the two groups, except DBP, which was higher in the control group prior to intervention. However, on regression analysis of all the probable confounding variables, none of the parameters seemed to affected the change in DBP that was observed.

#### During intervention

The key findings in BP at 20^th^ minute (during music) in group A were - SBP was similar to pre-intervention levels (indicating no effect /relaxation response), DBP increased significantly by 1.676 mm Hg, while HR insignificantly reduced. The various parasympathetic parameters of HRV [SDNN, RMSSD, VLF, HF (ms^2^)] reduced along with HR, TP (ms^2^) and LF (ms^2^). Mean NN interval increased, but not significantly. Thus a sympathetic predominance or reduction in parasympathetic activity was observed with music intervention. This might be due to the arousal effect of music as observed in few other studies (52–54).

In control group, SBP reduced very slightly, DBP increased, but was not significant, while HR continued to reduce throughout the 30 minutes’ duration. Adding to this finding, in control group sustained and significant increase in mean NN interval, along with parasympathetic HRV parameters along with TP and LF (ms^2^) was observed along with reduction in HR, implying increased parasympathetic activity, when a person is relaxing completely for 30 minutes’ duration, with larger amount of silence and natural sounds interspersing, for very short duration, in the mid 10 minutes (total 50 seconds). Nevertheless, the unit drop in HR was very less (hardly 1 beat per minute)

#### After intervention

After the intervention was stopped, in group A, SBP, HR increased mildly. The DBP increase was significant on comparison to pre-intervention levels (increase by 1.824 mm Hg). All the HRV parameters increased towards pre-intervention levels with reduction in HR (change being significant only for TP and LF). In the control group (group B), the SBP, DBP and HR reduced insignificantly after the intervention. Among the HRV parameters, mean NN interval, HR remained similar to during intervention levels. Other HRV parameters increased (change being significant for SDNN, TP and LF).

Note that LF power is produced by both SNS and PNS activity and is not a pure index of SNS drive. While SDNN, RMSSD, HF power is predominantly controlled by the PNS activity. Total power is said to reflect overall autonomic activity but has predominant vagal influence. (49).

#### Discussion of main findings

It can also be observed that passive listening task to *raga Bhimpalas* caused sympathetic arousal [as shown by increased DBP - indicating mild vasoconstriction in the periphery and drop in parasympathetic parameters of HRV – SDNN, RMSSD, HF (ms^2^)] during music, while regaining autonomic balance, after the music was stopped. This seems similar to the classic paper by Bernardi *et al*, where playing music for 2 minutes exhibited arousal response as against after stopping the music (32). Note that over ten minutes of music listening, the SBP did not change much, while HR reduced mildly, with increase in mean NN interval, though insignificant. The subjects involved in the current study were clinically normal, normotensives, (autonomically sound), and a large change in BP with music intervention, may be too high an expectation. In our previous study, music intervention caused a significant drop in DBP (∼2 mm Hg) among prehypertensives (41). In the current study, BP was measured acutely as subjects listened to music (in the lab) unlike in the previous study where BP was measured using 24 hour ambulatory BP device, prior to and after 3 months of intervention. Continuous BP monitoring, over 30 minutes, like ECG could have been better in indicating the real-time changes in BP. However, the observed significant DBP and HRV changes may be pointing towards the exciting / joyous emotion /arousal effect of *raga Bhimpalas*, as observed in the Indian Bollywood compositions based on this scale - *nainon mein badra chaaye (movie: mera saaya), E neele gagan ke tale* (*movie: Badshah*), *Kuch dil ne kaha* (*movie: Anupama*), *khilte hain gul yahan (movie: Sharmili)* (13).

This arousal response could be due to passive listening to an unfamiliar tune. Studies show that passive listening task produces a sympathetic response, when it is an emotionally arousing music (55). Similarly, in another study, authors interpreted that passive listening to positively valenced music increased HR, and listening to music in general was associated with a mind wandering state (56). As opposed to current study findings, Weiss *et al*., demonstrated greater pupil dilations (sympathetic arousal) during listening to familiar folk melody, compared to unfamiliar (novel) stimulus (57). Further in sports research, using music, it has been shown that unfamiliar relaxing music was the most relaxing (physiologically recorded using galvanic skin resistance, HR and peripheral temperature) and unfamiliar arousing music was the most arousing (58,59). A repeated exposure might have resulted in familiarity to the tune that was offered in this study and a different result. The probability of listening to music with an intention of relaxation, over a longer period of time, producing familiarity, under non-laboratory condition and thus a cumulative relaxation effect cannot be ruled out. To this thought, one study examined HRV among 13 students, after repetitive exposure to sedative music, excitatory music and no music condition. Each participant went through four sessions of one condition in a day. The LF and LF/HF ratio increased during sedative and excitatory music sessions but decreased during non-music conditions. The HF was higher during sedative music than during excitatory music listening but similar to non-music condition (60).

Non-expert listeners have a different physiological effect on listening to different genres of music (of different style and emotional outcome). In fifty untrained individuals, listening to atonal music was associated with a reduced HR and increased BP (SBP and DBP), possibly reflecting an increase in alertness and attention, psychological tension, and anxiety (61). Seventy percent of subjects in the current study were not trained in music. Further, regression analysis of the current study showed no effect of training on BP and HRV parameters.

The components in the music heard is also important to understand its physiological effect. Bowling et al., observed that melodies that are positive / excitatory had more major intervals (>200 cents) while negative/ subdued ragas have more minor intervals (6). A recent study on emotions caused by *thaats* (scales with all 7 notes) of Hindustani music by varying the tones, and tempo of Hindustani music concluded that *ragas* with major intervals (*shuddh swaras - shuddh Re and shuddh Ga*) were rated as ‘calm’ while those with minor intervals (*komal swaras - komal re and komal dha*) were rated as ‘sad’ (22). *Bhimpalas raga* is a unique scale with 3 perfect notes and equal number of major and minor notes. This might explain the mild sympathetic effect seen in the current study.

The frequency which is usually used to tune musical instruments is about 440 Hz. A study noted that 432 Hz music was associated with an insignificant decrease of SBP and DBP, a significant decrease in HR, compared to 440 Hz and that the subjects were more focused and more satisfied after they heard to music tuned at 432 Hz (62). In the present study music was played at Scale ‘E’ the frequency of which is 329.36 Hz – Note Sa (fundamental note).

The genre of music preferred by the subjects may be important as well. In the present study, experimenter chosen standardized music stimulus was used. Self-selected sedative music has been shown to induce both aroused and sedative emotions and a slight but significant increase in HR (63). In contrast another study, using sedative and stimulating music among cardiac rehabilitation patients, showed no effect of the type of music on BP (64). Music by Mozart, Strauss and the control group resulted in lower SBP, DBP; whereas listening to pop music (*ABBA*) caused no change. HR reduced significantly with Mozart music compared to control group. None of the above effects correlated with the music preference of the subjects (65).

In a crossover study of 40 min of active and passive intervention, active music therapy reduced LF/HF while passive music intervention (as seen in present study) increased LF/HF (25). Acute effect of different genres of music (heavy metal music, classical baroque music) on HRV parameters showed that listening to heavy metal music reduced SDNN significantly. LF (ms^2^ and nu) reduced with both types of music, while HF (ms^2^) reduced only with heavy metal music and LF/HF ratio reduced with classical baroque music. Authors thus concluded that heavy metal music decreased the autonomic modulation, while exposure to a classical baroque music reduced sympathetic regulation on the heart (66,67). When HRV was analysed during no sound condition RMSSD significantly increased, which authors attributed to the supine posture, when cardiac PNS inputs were maximal (68). In the current study protocol supine posture was followed for all participants, with the control group rested for complete 30 minutes, without any disturbance, while music intervention group, rested with music being played in the middle 10 minutes. Music chosen was not guided with breathing frequency, though the respiratory rate was recorded. Music combined with guided breathing exercises have shown better control of physiological parameters in a few studies (69,70). Nevertheless, recently a study concluded that both listening to music and deep breathing exercise were associated with a clinically significant reduction in SBP and DBP and that deep breathing exercise did not augment the benefit of music in reducing BP (71). A review based on the effect of auditory stimulation and cardiac autonomic regulation hypothesized that dopamine release in the striatal system, that is induced by pleasurable songs was involved in cardiac autonomic regulation (72).

The strengths of the study are that, to the best of our knowledge, for the first time an Indian melodic scale has been studied systematically and scientifically, via a randomized trial (avoiding different types of bias), among normal healthy individuals, with recording of various physiological parameters. All parameters were free from measurement errors as the recordings were performed by a single, well trained, but blinded, research assistant (reducing observer’s error) along with validation from the PI (who was also blinded) for accuracy of the data collected. Further the devices used to collect the data was standard, reliable and well validated, through prior research studies. The chosen music intervention was standardized and was based on existing music literatures. Music used was composed of pure *alaap* and drone instrument. Percussion instruments and lyrical component was avoided, as they have their own respective effects on physiological parameters. Passive listening to music was chosen to maintain uniformity, as active music making may not be an option for all subjects. The control group was well matched and the intervention received by the control group was also standardized. The sample size was calculated based on prior research work, with appropriate power and was adequate to show measurable, significant effects of the intervention. Subjects of both genders, with homogenous age groups were compared. Thus the evidence obtained through this work is true and scientifically authentic.

The limitations of the study were that choice of music was not given to the subjects. Literature review shows music has better effect, especially with respect to pain, when self-chosen (73–77). However there have been quite a number of other research works that prove that experimenter chosen music is better than self-chosen music (13) and a few others showing that choice of music did not matter (78). One fact of this study was that the participants knew the aim of the study, and might have listened to the music with a particular intention, that is different from that of control group, as once the intervention started, control group knew they were not in the music intervention group. This limitation was difficult to overcome, as research question demanded this design and the results obtained may be interpreted with this background. A subject’s involvement in the music and subjective emotions were not captured. Self-reported parameters are however less reliable than actual physiological recordings. Though all subjects were clinically normal, laboratory measurements of their blood/serum or urine was not conducted to conclusively say that everyone was normal. Further extraction of the musical components through pre-determined software, and its correlation with the continuous monitoring of BP and HRV may further elaborate on the cause-effect response seen.

## Conclusion

For the first time, through this study we have shown the acute effect of Indian music on cardiovascular electrophysiological parameters, in a systematic fashion. Passive listening to a north Indian Hindustani classical musical scale, *raga bhimpalas* caused mild arousal response during intervention, that trended to return to baseline levels after the intervention was stopped. This may be attributed to a combination of major and minor intervals in the scale as well as a normal response to an unfamiliar stimulus, that was heard with enough attention so as to produce a mild sympathetic arousal. Future studies should try to evaluate the physiological responses during passive listening to different genres, scales of music, and the musical, after familiarizing to the stimuli.

## Credit roles

Conceptualization, Funding acquisition – UKK; Data curation – UKK, VJ, Formal analysis – UKK, RK, NSM; Investigation – UKK, VJ; Methodology – UKK, VJ, GJ; Project administration – UKK, VJ, GJ, NSM, RK; Resources – UKK, VJ; Software – UKK, RK, NSM; Supervision – UKK, VJ, GJ, VSP; Validation – RK, NSM; Writing original draft – UKK, GJ; Review & editing – VJ, RK, NSM, VSP.

## Acknowledgement

Authors acknowledge funding agencies, administrative support received at Ramaiah Medical college, Bangalore. We acknowledge the exclusive recording shared by *Vidhwan Pandit Pravin Godhkhindi*, an eminent flautist, for this study.

## Footnotes

1 The repeated measures ANOVA tests for whether there are any differences between related population means. The null hypothesis (H_0_) states that the means are equal: *H*_*0*_: *µ*_*1*_ *= µ*_*2*_ *= µ*_*3*_ *= … = µ*_*k*_ where *µ* = population mean and *k* = number of related groups. The alternative hypothesis (H_A_) states that the related population means are not equal (at least one mean is different to another mean): *H*_*A*_: *at least two means are significantly different*

