## Supplemental file for "Acute effects of passive listening to Indian musical scale on blood pressure and heart rate variability among healthy young individuals – a randomized controlled trial"

**Table 1: Pairwise comparison of pre (1), during (2) and post intervention (3) SBP between the groups A (music intervention) and group B (control group) (N=34 in each group)**

| RANDOM GROUP | | | Mean Difference (I-J) | Std. Error | Sig.^a^ | 95% Confidence Interval for Difference^a^ | |
| --- | --- | --- | --- | --- | --- | --- | --- |
|  |  |  |  |  |  | Lower Bound | Upper Bound |
| A | 1 | 2 | .206 | .890 | 1.000 | -2.040 | 2.452 |
|  |  | 3 | -1.794 | .869 | .141 | -3.986 | .398 |
|  | 2 | 1 | -.206 | .890 | 1.000 | -2.452 | 2.040 |
|  |  | 3 | -2.000 | .910 | .106 | -4.296 | .296 |
|  | 3 | 1 | 1.794 | .869 | .141 | -.398 | 3.986 |
|  |  | 2 | 2.000 | .910 | .106 | -.296 | 4.296 |
| B | 1 | 2 | .353 | .690 | 1.000 | -1.387 | 2.093 |
|  |  | 3 | 1.941 | 1.349 | .479 | -1.461 | 5.344 |
|  | 2 | 1 | -.353 | .690 | 1.000 | -2.093 | 1.387 |
|  |  | 3 | 1.588 | 1.336 | .729 | -1.781 | 4.958 |
|  | 3 | 1 | -1.941 | 1.349 | .479 | -5.344 | 1.461 |
|  |  | 2 | -1.588 | 1.336 | .729 | -4.958 | 1.781 |
| Based on estimated marginal means | | | | | | | |
| a. Adjustment for multiple comparisons: Bonferroni. | | | | | | | |

**Table 2: Pairwise comparison of pre (1), during (2) and post intervention (3) DBP between the groups A (music intervention) and group B (control group)**

| RANDOM GROUP | | | Mean Difference (I-J) | Std. Error | Sig.^b^ | 95% Confidence Interval for Difference^b^ | |
| --- | --- | --- | --- | --- | --- | --- | --- |
|  |  |  |  |  |  | Lower Bound | Upper Bound |
| A | 1 | 2 | -1.676^*^ | .540 | **0.012** | -3.039 | -.314 |
|  |  | 3 | -1.824^*^ | .653 | **0.026** | -3.471 | -.176 |
|  | 2 | 1 | 1.676^*^ | .540 | .012 | .314 | 3.039 |
|  |  | 3 | -.147 | .696 | 1.000 | -1.904 | 1.610 |
|  | 3 | 1 | 1.824^*^ | .653 | 0.026 | .176 | 3.471 |
|  |  | 2 | .147 | .696 | 1.000 | -1.610 | 1.904 |
| B | 1 | 2 | -1.235 | .938 | .591 | -3.602 | 1.132 |
|  |  | 3 | -.618 | 1.155 | 1.000 | -3.530 | 2.295 |
|  | 2 | 1 | 1.235 | .938 | .591 | -1.132 | 3.602 |
|  |  | 3 | .618 | .677 | 1.000 | -1.090 | 2.325 |
|  | 3 | 1 | .618 | 1.155 | 1.000 | -2.295 | 3.530 |
|  |  | 2 | -.618 | .677 | 1.000 | -2.325 | 1.090 |
| Based on estimated marginal means | | | | | | | |
| *. The mean difference is significant at the .05 level. | | | | | | | |
| b. Adjustment for multiple comparisons: Bonferroni. | | | | | | | |

**Table 3: Pairwise comparison of pre (1), during (2) and post intervention (3) HR between the groups A (music intervention) and group B (control group) (N=34 in each group)**

| RANDOM GROUP | | | Mean Difference (I-J) | Std. Error | Sig.^a^ | 95% Confidence Interval for Difference^a^ | |
| --- | --- | --- | --- | --- | --- | --- | --- |
|  |  |  |  |  |  | Lower Bound | Upper Bound |
| A | 1 | 2 | .235 | 1.070 | 1.000 | -2.463 | 2.934 |
|  |  | 3 | -.118 | .917 | 1.000 | -2.431 | 2.195 |
|  | 2 | 1 | -.235 | 1.070 | 1.000 | -2.934 | 2.463 |
|  |  | 3 | -.353 | .990 | 1.000 | -2.850 | 2.144 |
|  | 3 | 1 | .118 | .917 | 1.000 | -2.195 | 2.431 |
|  |  | 2 | .353 | .990 | 1.000 | -2.144 | 2.850 |
| B | 1 | 2 | .412 | 1.626 | 1.000 | -3.689 | 4.513 |
|  |  | 3 | .971 | 1.589 | 1.000 | -3.037 | 4.978 |
|  | 2 | 1 | -.412 | 1.626 | 1.000 | -4.513 | 3.689 |
|  |  | 3 | .559 | .989 | 1.000 | -1.937 | 3.054 |
|  | 3 | 1 | -.971 | 1.589 | 1.000 | -4.978 | 3.037 |
|  |  | 2 | -.559 | .989 | 1.000 | -3.054 | 1.937 |
| Based on estimated marginal means | | | | | | | |
| a. Adjustment for multiple comparisons: Bonferroni. | | | | | | | |

**Table 4: Comparison of pre, during and post intervention HRV (Absolute values) between the groups (N=34 in each group)**

|  | **Group A** | | | **Group B** | | |
| --- | --- | --- | --- | --- | --- | --- |
|  | **Pre** | **During** | **Post** | **Pre** | **During** | **Post** |
| **Mean NN (ms)** | 830.7, 78.99 | 840.1, 83.24 | 844.9, 83 | 843.9, 131.27 | 862.7, 131.19 | 863.4, 128.42 |
| **HR (bpm)** | 72.9, 7.09 | 72.1, 7.26 | 71.7, 7.18 | 72.8, 11.29 | 71.1, 10.49 | 71.0, 10.59 |
| **SDNN** | 61.2, 24.57 | 56.8, 22.09 | 60.2, 24.93 | 62.1, 24.43 | 65.0, 24.07 | 71.6, 27.21 |
| **RMSSD** | 57.6, 32.69 | 55.4, 31.55 | 57.7, 31.04 | 59.0, 31.95 | 62.6, 31.54 | 66.9, 33.67 |
| **NN50** | 267.2, 229.95 | 223.0, 161.85 | 226.9, 172.94 | 236.5, 183.12 | 246.9, 162.02 | 247.1, 137.21 |
| **pNN50** | 30.4, 21.83 | 30.7, 24.11 | 31.5, 22.51 | 31.6, 22.52 | 33.5, 21.85 | 35.4, 21.06 |
| **TP (ms^2^)** | 4060.0, 3571.09 | 3471.0, 2862.87 | 4246.3, 4013.90 | 4177.4, 3342.32 | 4567.3, 3465.36 | 5255.5, 3774.23 |
| **VLF (ms^2^)** | 1206.9, 915.00 | 977.4, 686.24 | 1271.2, 1994.21 | 1151.5, 1038.37 | 1336.5, 1264.74 | 1404.8, 1012.25 |
| **LF (ms^2^)** | 965.7, 980.02 | 811.5, 703.91 | 1058.1, 943.60 | 1171.4, 1188.56 | 1296.0, 1266.64 | 1539.9, 1426.75 |
| **LF (nu)** | 38.5, 14.84 | 38.4, 16.99 | 41.8, 15.87 | 41.7, 13.86 | 41.4, 15.37 | 43.5, 15.09 |
| **HF (ms^2^)** | 1620.3, 1902.51 | 1472.0, 1737.08 | 1672.4, 1996.48 | 1591.1, 1544.96 | 1709.7, 1479.91 | 2047.8, 1897.57 |
| **HF (nu)** | 52.0, 15.01 | 53.3, 17.17 | 49.4, 15.15 | 49.4, 14.23 | 51.4, 14.08 | 49.4, 14.05 |
| **LF/HF** | 0.9, 0.69 | 1.0, 0.76 | 1.0, 0.71 | 1.1, 0.74 | 1.0, 0.84 | 1.1, 0.77 |

**Table 5 (a to m): Pairwise comparison of pre (1), during (2) and post intervention (3) HRV between the groups A (music intervention) and group B (control group) (N=34 in each group)**

| a) Pairwise Comparisons Mean NN | | | | | | | |
| --- | --- | --- | --- | --- | --- | --- | --- |
| RANDOM GROUP | | | Mean Difference (I-J) | Std. Error | Sig.^a^ | 95% Confidence Interval for Difference^a^ | |
|  |  |  |  |  |  | Lower Bound | Upper Bound |
| C | 1 | 2 | -.004 | .003 | .433 | -.011 | .003 |
|  |  | 3 | -.006 | .003 | .171 | -.014 | .002 |
|  | 2 | 1 | .004 | .003 | .433 | -.003 | .011 |
|  |  | 3 | -.002 | .003 | 1.000 | -.009 | .004 |
|  | 3 | 1 | .006 | .003 | .171 | -.002 | .014 |
|  |  | 2 | .002 | .003 | 1.000 | -.004 | .009 |
| D | 1 | 2 | -.009^*^ | .002 | .005 | -.015 | -.002 |
|  |  | 3 | -.009 | .005 | .205 | -.021 | .003 |
|  | 2 | 1 | .009^*^ | .002 | .005 | .002 | .015 |
|  |  | 3 | .000 | .003 | 1.000 | -.008 | .007 |
|  | 3 | 1 | .009 | .005 | .205 | -.003 | .021 |
|  |  | 2 | .000 | .003 | 1.000 | -.007 | .008 |
| Based on estimated marginal means | | | | | | | |
| *. The mean difference is significant at the .05 level. | | | | | | | |
| a. Adjustment for multiple comparisons: Bonferroni. | | | | | | | |

| b) Pairwise Comparisons of HR | | | | | | | |
| --- | --- | --- | --- | --- | --- | --- | --- |
| Measure: MEASURE_1 | | | | | | | |
| RANDOM GROUP | | | Mean Difference (I-J) | Std. Error | Sig.^a^ | 95% Confidence Interval for Difference^a^ | |
|  |  |  |  |  |  | Lower Bound | Upper Bound |
| C | 1 | 2 | .782 | .505 | .392 | -.491 | 2.055 |
|  |  | 3 | 1.200 | .564 | .123 | -.223 | 2.622 |
|  | 2 | 1 | -.782 | .505 | .392 | -2.055 | .491 |
|  |  | 3 | .417 | .437 | 1.000 | -.685 | 1.520 |
|  | 3 | 1 | -1.200 | .564 | .123 | -2.622 | .223 |
|  |  | 2 | -.417 | .437 | 1.000 | -1.520 | .685 |
| D | 1 | 2 | 1.669^*^ | .441 | .002 | .557 | 2.781 |
|  |  | 3 | 1.799 | .853 | .128 | -.354 | 3.951 |
|  | 2 | 1 | -1.669^*^ | .441 | .002 | -2.781 | -.557 |
|  |  | 3 | .130 | .518 | 1.000 | -1.176 | 1.436 |
|  | 3 | 1 | -1.799 | .853 | .128 | -3.951 | .354 |
|  |  | 2 | -.130 | .518 | 1.000 | -1.436 | 1.176 |
| Based on estimated marginal means | | | | | | | |
| *. The mean difference is significant at the .05 level. | | | | | | | |
| a. Adjustment for multiple comparisons: Bonferroni. | | | | | | | |

| c) Pairwise Comparisons of SDNN | | | | | | | |
| --- | --- | --- | --- | --- | --- | --- | --- |
| RANDOM GROUP | | | Mean Difference (I-J) | Std. Error | Sig.^a^ | 95% Confidence Interval for Difference^a^ | |
|  |  |  |  |  |  | Lower Bound | Upper Bound |
| C | 1 | 2 | .027 | .015 | .214 | -.010 | .064 |
|  |  | 3 | .033 | .031 | .895 | -.046 | .112 |
|  | 2 | 1 | -.027 | .015 | .214 | -.064 | .010 |
|  |  | 3 | .006 | .034 | 1.000 | -.079 | .091 |
|  | 3 | 1 | -.033 | .031 | .895 | -.112 | .046 |
|  |  | 2 | -.006 | .034 | 1.000 | -.091 | .079 |
| D | 1 | 2 | -.020 | .014 | .453 | -.055 | .015 |
|  |  | 3 | -.054^*^ | .018 | .014 | -.099 | -.009 |
|  | 2 | 1 | .020 | .014 | .453 | -.015 | .055 |
|  |  | 3 | -.034^*^ | .012 | .024 | -.064 | -.004 |
|  | 3 | 1 | .054^*^ | .018 | .014 | .009 | .099 |
|  |  | 2 | .034^*^ | .012 | .024 | .004 | .064 |
| Based on estimated marginal means | | | | | | | |
| *. The mean difference is significant at the .05 level. | | | | | | | |
| a. Adjustment for multiple comparisons: Bonferroni. | | | | | | | |

| d) Pairwise Comparisons of RMSSD | | | | | | | |
| --- | --- | --- | --- | --- | --- | --- | --- |
| RANDOM GROUP | | | Mean Difference (I-J) | Std. Error | Sig.^a^ | 95% Confidence Interval for Difference^a^ | |
|  |  |  |  |  |  | Lower Bound | Upper Bound |
| C | 1 | 2 | .016 | .018 | 1.000 | -.029 | .061 |
|  |  | 3 | -.008 | .015 | 1.000 | -.047 | .031 |
|  | 2 | 1 | -.016 | .018 | 1.000 | -.061 | .029 |
|  |  | 3 | -.024 | .016 | .442 | -.065 | .017 |
|  | 3 | 1 | .008 | .015 | 1.000 | -.031 | .047 |
|  |  | 2 | .024 | .016 | .442 | -.017 | .065 |
| D | 1 | 2 | -.059 | .032 | .234 | -.140 | .023 |
|  |  | 3 | -.082 | .037 | .094 | -.175 | .010 |
|  | 2 | 1 | .059 | .032 | .234 | -.023 | .140 |
|  |  | 3 | -.023 | .012 | .179 | -.054 | .007 |
|  | 3 | 1 | .082 | .037 | .094 | -.010 | .175 |
|  |  | 2 | .023 | .012 | .179 | -.007 | .054 |
| Based on estimated marginal means | | | | | | | |
| a. Adjustment for multiple comparisons: Bonferroni. | | | | | | | |

| e) Pairwise Comparisons of NN50 | | | | | | | |
| --- | --- | --- | --- | --- | --- | --- | --- |
| RANDOM GROUP | | | Mean Difference (I-J) | Std. Error | Sig.^a^ | 95% Confidence Interval for Difference^a^ | |
|  |  |  |  |  |  | Lower Bound | Upper Bound |
| C | 1 | 2 | .025 | .044 | 1.000 | -.087 | .137 |
|  |  | 3 | .004 | .042 | 1.000 | -.102 | .110 |
|  | 2 | 1 | -.025 | .044 | 1.000 | -.137 | .087 |
|  |  | 3 | -.021 | .033 | 1.000 | -.103 | .062 |
|  | 3 | 1 | -.004 | .042 | 1.000 | -.110 | .102 |
|  |  | 2 | .021 | .033 | 1.000 | -.062 | .103 |
| D | 1 | 2 | -.064 | .042 | .419 | -.170 | .042 |
|  |  | 3 | -.067 | .059 | .792 | -.217 | .082 |
|  | 2 | 1 | .064 | .042 | .419 | -.042 | .170 |
|  |  | 3 | -.004 | .037 | 1.000 | -.098 | .091 |
|  | 3 | 1 | .067 | .059 | .792 | -.082 | .217 |
|  |  | 2 | .004 | .037 | 1.000 | -.091 | .098 |
| Based on estimated marginal means | | | | | | | |
| a. Adjustment for multiple comparisons: Bonferroni. | | | | | | | |

| f) Pairwise Comparisons of pNN50 | | | | | | | |
| --- | --- | --- | --- | --- | --- | --- | --- |
| RANDOM GROUP | | | Mean Difference (I-J) | Std. Error | Sig.^a^ | 95% Confidence Interval for Difference^a^ | |
|  |  |  |  |  |  | Lower Bound | Upper Bound |
| C | 1 | 2 | .002 | .037 | 1.000 | -.091 | .095 |
|  |  | 3 | -.019 | .049 | 1.000 | -.141 | .103 |
|  | 2 | 1 | -.002 | .037 | 1.000 | -.095 | .091 |
|  |  | 3 | -.021 | .042 | 1.000 | -.126 | .085 |
|  | 3 | 1 | .019 | .049 | 1.000 | -.103 | .141 |
|  |  | 2 | .021 | .042 | 1.000 | -.085 | .126 |
| D | 1 | 2 | -.065 | .040 | .347 | -.166 | .037 |
|  |  | 3 | -.102 | .063 | .351 | -.263 | .058 |
|  | 2 | 1 | .065 | .040 | .347 | -.037 | .166 |
|  |  | 3 | -.037 | .035 | .864 | -.125 | .050 |
|  | 3 | 1 | .102 | .063 | .351 | -.058 | .263 |
|  |  | 2 | .037 | .035 | .864 | -.050 | .125 |
| Based on estimated marginal means | | | | | | | |
| a. Adjustment for multiple comparisons: Bonferroni. | | | | | | | |

| g) Pairwise Comparisons of TP ms2 | | | | | | | |
| --- | --- | --- | --- | --- | --- | --- | --- |
| RANDOM GROUP | | | Mean Difference (I-J) | Std. Error | Sig.^a^ | 95% Confidence Interval for Difference^a^ | |
|  |  |  |  |  |  | Lower Bound | Upper Bound |
| C | 1 | 2 | .058 | .029 | .146 | -.014 | .130 |
|  |  | 3 | -.011 | .022 | 1.000 | -.067 | .044 |
|  | 2 | 1 | -.058 | .029 | .146 | -.130 | .014 |
|  |  | 3 | -.070 | .031 | .100 | -.149 | .010 |
|  | 3 | 1 | .011 | .022 | 1.000 | -.044 | .067 |
|  |  | 2 | .070 | .031 | .100 | -.010 | .149 |
| D | 1 | 2 | -.057 | .033 | .293 | -.141 | .027 |
|  |  | 3 | -.133^*^ | .048 | .026 | -.253 | -.013 |
|  | 2 | 1 | .057 | .033 | .293 | -.027 | .141 |
|  |  | 3 | -.076 | .036 | .124 | -.166 | .014 |
|  | 3 | 1 | .133^*^ | .048 | .026 | .013 | .253 |
|  |  | 2 | .076 | .036 | .124 | -.014 | .166 |
| Based on estimated marginal means | | | | | | | |
| *. The mean difference is significant at the .05 level. | | | | | | | |
| a. Adjustment for multiple comparisons: Bonferroni. | | | | | | | |

| h) Pairwise Comparisons of VLF ms2 | | | | | | | |
| --- | --- | --- | --- | --- | --- | --- | --- |
| RANDOM GROUP | | | Mean Difference (I-J) | Std. Error | Sig.^a^ | 95% Confidence Interval for Difference^a^ | |
|  |  |  |  |  |  | Lower Bound | Upper Bound |
| C | 1 | 2 | .074 | .042 | .257 | -.031 | .179 |
|  |  | 3 | .081 | .048 | .311 | -.041 | .202 |
|  | 2 | 1 | -.074 | .042 | .257 | -.179 | .031 |
|  |  | 3 | .007 | .051 | 1.000 | -.121 | .135 |
|  | 3 | 1 | -.081 | .048 | .311 | -.202 | .041 |
|  |  | 2 | -.007 | .051 | 1.000 | -.135 | .121 |
| D | 1 | 2 | -.093 | .059 | .379 | -.243 | .057 |
|  |  | 3 | -.113 | .060 | .203 | -.263 | .038 |
|  | 2 | 1 | .093 | .059 | .379 | -.057 | .243 |
|  |  | 3 | -.020 | .051 | 1.000 | -.148 | .108 |
|  | 3 | 1 | .113 | .060 | .203 | -.038 | .263 |
|  |  | 2 | .020 | .051 | 1.000 | -.108 | .148 |
| Based on estimated marginal means | | | | | | | |
| a. Adjustment for multiple comparisons: Bonferroni. | | | | | | | |

| i) Pairwise Comparisons of LF ms2 | | | | | | | |
| --- | --- | --- | --- | --- | --- | --- | --- |
| RANDOM GROUP | | | Mean Difference (I-J) | Std. Error | Sig.^b^ | 95% Confidence Interval for Difference^b^ | |
|  |  |  |  |  |  | Lower Bound | Upper Bound |
| C | 1 | 2 | .090 | .046 | .181 | -.027 | .207 |
|  |  | 3 | -.035 | .043 | 1.000 | -.144 | .074 |
|  | 2 | 1 | -.090 | .046 | .181 | -.207 | .027 |
|  |  | 3 | -.125^*^ | .037 | .005 | -.218 | -.032 |
|  | 3 | 1 | .035 | .043 | 1.000 | -.074 | .144 |
|  |  | 2 | .125^*^ | .037 | .005 | .032 | .218 |
| D | 1 | 2 | -.056 | .043 | .624 | -.165 | .054 |
|  |  | 3 | -.127 | .052 | .059 | -.258 | .004 |
|  | 2 | 1 | .056 | .043 | .624 | -.054 | .165 |
|  |  | 3 | -.072^*^ | .028 | .049 | -.143 | .000 |
|  | 3 | 1 | .127 | .052 | .059 | -.004 | .258 |
|  |  | 2 | .072^*^ | .028 | .049 | .000 | .143 |
| Based on estimated marginal means | | | | | | | |
| *. The mean difference is significant at the .05 level. | | | | | | | |
| b. Adjustment for multiple comparisons: Bonferroni. | | | | | | | |

| j) Pairwise Comparisons of LF nu | | | | | | | |
| --- | --- | --- | --- | --- | --- | --- | --- |
| RANDOM GROUP | | | Mean Difference (I-J) | Std. Error | Sig.^a^ | 95% Confidence Interval for Difference^a^ | |
|  |  |  |  |  |  | Lower Bound | Upper Bound |
| C | 1 | 2 | .044 | .037 | .711 | -.049 | .138 |
|  |  | 3 | -.002 | .038 | 1.000 | -.097 | .093 |
|  | 2 | 1 | -.044 | .037 | .711 | -.138 | .049 |
|  |  | 3 | -.047 | .019 | .062 | -.095 | .002 |
|  | 3 | 1 | .002 | .038 | 1.000 | -.093 | .097 |
|  |  | 2 | .047 | .019 | .062 | -.002 | .095 |
| D | 1 | 2 | .004 | .016 | 1.000 | -.038 | .045 |
|  |  | 3 | -.014 | .021 | 1.000 | -.067 | .038 |
|  | 2 | 1 | -.004 | .016 | 1.000 | -.045 | .038 |
|  |  | 3 | -.018 | .016 | .788 | -.057 | .022 |
|  | 3 | 1 | .014 | .021 | 1.000 | -.038 | .067 |
|  |  | 2 | .018 | .016 | .788 | -.022 | .057 |
| Based on estimated marginal means | | | | | | | |
| a. Adjustment for multiple comparisons: Bonferroni. | | | | | | | |

| k) Pairwise Comparisons of HF ms2 | | | | | | | |
| --- | --- | --- | --- | --- | --- | --- | --- |
| RANDOM GROUP | | | Mean Difference (I-J) | Std. Error | Sig.a | 95% Confidence Interval for Differencea | |
|  |  |  |  |  |  | Lower Bound | Upper Bound |
| C | 1 | 2 | .041 | .035 | .759 | -.048 | .130 |
|  |  | 3 | .018 | .035 | 1.000 | -.071 | .106 |
|  | 2 | 1 | -.041 | .035 | .759 | -.130 | .048 |
|  |  | 3 | -.023 | .043 | 1.000 | -.131 | .084 |
|  | 3 | 1 | -.018 | .035 | 1.000 | -.106 | .071 |
|  |  | 2 | .023 | .043 | 1.000 | -.084 | .131 |
| D | 1 | 2 | -.097 | .046 | .123 | -.213 | .018 |
|  |  | 3 | -.140 | .058 | **.066** | -.286 | .007 |
|  | 2 | 1 | .097 | .046 | .123 | -.018 | .213 |
|  |  | 3 | -.042 | .027 | .372 | -.110 | .025 |
|  | 3 | 1 | .140 | .058 | .066 | -.007 | .286 |
|  |  | 2 | .042 | .027 | .372 | -.025 | .110 |
| Based on estimated marginal means | | | | | | | |
| a. Adjustment for multiple comparisons: Bonferroni. | | | | | | | |

| l) Pairwise Comparisons of HF nu | | | | | | | |
| --- | --- | --- | --- | --- | --- | --- | --- |
| RANDOM GROUP | | | Mean Difference (I-J) | Std. Error | Sig.^a^ | 95% Confidence Interval for Difference^a^ | |
|  |  |  |  |  |  | Lower Bound | Upper Bound |
| C | 1 | 2 | -.005 | .017 | 1.000 | -.048 | .039 |
|  |  | 3 | .024 | .017 | .521 | -.020 | .068 |
|  | 2 | 1 | .005 | .017 | 1.000 | -.039 | .048 |
|  |  | 3 | .029 | .019 | .430 | -.020 | .077 |
|  | 3 | 1 | -.024 | .017 | .521 | -.068 | .020 |
|  |  | 2 | -.029 | .019 | .430 | -.077 | .020 |
| D | 1 | 2 | -.038 | .035 | .852 | -.126 | .050 |
|  |  | 3 | -.028 | .037 | 1.000 | -.122 | .065 |
|  | 2 | 1 | .038 | .035 | .852 | -.050 | .126 |
|  |  | 3 | .010 | .014 | 1.000 | -.025 | .045 |
|  | 3 | 1 | .028 | .037 | 1.000 | -.065 | .122 |
|  |  | 2 | -.010 | .014 | 1.000 | -.045 | .025 |
| Based on estimated marginal means | | | | | | | |
| a. Adjustment for multiple comparisons: Bonferroni. | | | | | | | |

| m) Pairwise Comparisons of LF/HF | | | | | | | |
| --- | --- | --- | --- | --- | --- | --- | --- |
| RANDOM GROUP | | | Mean Difference (I-J) | Std. Error | Sig.^a^ | 95% Confidence Interval for Difference^a^ | |
|  |  |  |  |  |  | Lower Bound | Upper Bound |
| C | 1 | 2 | .002 | .038 | 1.000 | -.094 | .097 |
|  |  | 3 | -.060 | .041 | .453 | -.164 | .043 |
|  | 2 | 1 | -.002 | .038 | 1.000 | -.097 | .094 |
|  |  | 3 | -.062 | .039 | .356 | -.159 | .036 |
|  | 3 | 1 | .060 | .041 | .453 | -.043 | .164 |
|  |  | 2 | .062 | .039 | .356 | -.036 | .159 |
| D | 1 | 2 | .002 | .043 | 1.000 | -.106 | .110 |
|  |  | 3 | -.028 | .051 | 1.000 | -.158 | .102 |
|  | 2 | 1 | -.002 | .043 | 1.000 | -.110 | .106 |
|  |  | 3 | -.030 | .031 | 1.000 | -.108 | .049 |
|  | 3 | 1 | .028 | .051 | 1.000 | -.102 | .158 |
|  |  | 2 | .030 | .031 | 1.000 | -.049 | .108 |
| Based on estimated marginal means | | | | | | | |
| a. Adjustment for multiple comparisons: Bonferroni. | | | | | | | |

**Figure 1: Scale of Bhimpalasi raga notes as represented in Indian music on western scale** (1,2)**:**

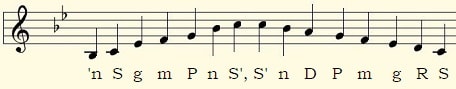

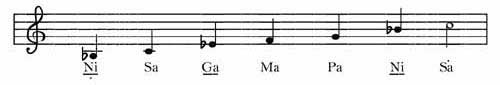

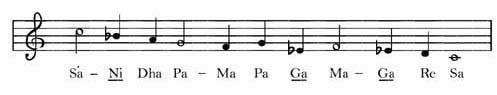

It is a pentatonic ascent and heptatonic descent.

**Table 6: Names & equivalents of the 12 basic notes of Indian music**

| **Carnatic** | | | | | **Western** | | | **Hindustani** | | |
| --- | --- | --- | --- | --- | --- | --- | --- | --- | --- | --- |
| ***Full Name*** | | ***Short forms*** | | ***Alt^1^*** | ***Interval*** | ***Cents^2^*** | ***Note^3^*** | ***Full Name*** | ***Short forms*** | |
| 1 | Shadjam | Sa | S |  | P1 | 0 | C | Shadj | S | S |
| 2 | Suddha Rishabham | Ri1 | R1 |  | m2 | 100 | D♭ | Komal Rishab | kR | R1 |
| 3 | Chatusruthi Rishabham | Ri2 | R2 |  | M2 | 200 | D | Suddh Rishab | R | R2 |
|  | Suddha Gantharam**^4^** | Ga1 | G1 | G0 |  |  | E♭♭ |  |  |  |
| 4 | Shatsruthi Rishabham**^4^** | Ri3 | R3 |  | m3 | 300 | D*#* | Komal Gandhar | kG | G1 |
|  | Saadarana Gantharam | Ga2 | G2 | G1 |  |  | E♭ |  |  |  |
| 5 | Antara Gantharam | Ga3 | G3 | G2 | M3 | 400 | E | Suddh Gandhar | G | G2 |
| 6 | Suddha Madhyamam | Ma1 | M1 |  | P4 | 500 | F | Suddh Madhyam | M | M1 |
| 7 | Prati Madhyamam | Ma2 | M2 |  | 4 | 600 | F*#* | Tivra Madhyam | tM | M2 |
| 8 | Panchamam | Pa | P |  | P5 | 700 | G | Pancham | P | P |
| 9 | Suddha Dhaivatham | Da1 | D1 |  | m6 | 800 | A♭ | Komal Dhaivat | kD | D1 |
| 10 | Chatusruthi Dhaivatham | Da2 | D2 |  | M6 | 900 | A | Suddh Dhaivat | D | D2 |
|  | Suddha Nishadham**^4^** | Ni1 | N1 | N0 |  |  | B♭♭ |  |  |  |
| 11 | Shatsruthi Dhaivatham**^4^** | Da3 | D3 |  | m7 | 1000 | A*#* | Komal Nishad | kN | N1 |
|  | Kaisiki Nishadham | Ni2 | N2 | N1 |  |  | B♭ |  |  |  |
| 12 | Kaakali Nishadham | Ni3 | N3 | N2 | M7 | 1100 | B | Suddh Nishad | N | N2 |

**^1^** Instead of labeling the 3 Ga-s and the 3 Ni-s using {1,2,3}, some authors have used {0,1,2} instead. This alternate numbering scheme also makes comparison to the Western and Hindustani notes easier. Both conventions are present in the literature, sometimes causing unnecessary confusion to novices! Note also that in this alternate scheme, all "vivadhi notes" are labeled by 0 or 3, and all notes labeled with a 0 or 3 are vivadhi.
**^2^** Western Equal Temperament tuning.
**^3^** Note names if and only if "C" is chosen as the tonic. If another note, say F#, is chosen as the tonic, then this column would have to be modified. The point is that the Indian notes correspond to the intervals - P1, m2, M2 , ..., M7 - rather than to absolute pitches/frequencies. The artist can choose Sa/the tonic to be any frequency he or she desires at the beginning of a performance.
**^4^** Vivadhi notes. These are are not new intervals/swarasthanams, but are enharmonic to 4 other "normal" notes. Ie, out of the sixteen Carnatic notes listed, there are only twelve unique intervals.

**Table 7: Svara and their names in North Indian system of raga and its equivalent western scale notes (3)**

| Svara in North Indian system of *raga* | | | | | | | |
| --- | --- | --- | --- | --- | --- | --- | --- |
| Svara (Long) | Ṣaḍja | Ṛiṣabha | Gāndhāra | Madhyama | Pañcama | Dhaivata | Niṣāda |
| Svara (Short) | Sa | Re | Ga | Ma | Pa | Dha | Ni |
| 12 Varieties (names) | C (shadja) | D♭ (komal re) D (shuddha re) | E♭ (komal ga) E (shuddha ga) | F (shuddha ma) F♯ (teevra ma) | G (panchama) | A♭ (komal dha) A (shuddha dha) | B♭ (komal ni) B (shuddha ni) |

**Just intonation and 12 - equal temperament:**

Just intonation or pure intonation is the tuning of musical intervals as whole number ratios (such as 3:2 or 4:3) of frequencies. Any interval tuned in this way is called a just interval. Just intervals (and chords created by combining them) consist of members of a single harmonic series of a (lower) implied fundamental. For example, in the diagram at right, the notes G and middle C (labeled 3 and 4), are both members of the harmonic series of the lowest C and their frequencies will be 3 and 4 times, respectively, the fundamental frequency; thus, their interval ratio will be 4:3. If the frequency of the fundamental is 64 Hertz, the frequencies of the two notes in question would be 192 and 256. Instruments are not always tuned using these intervals.

In the Western world, instruments of fixed pitch, such as pianos, are typically tuned using equal temperament, in which intervals other than octaves consist of irrational-number frequency ratios. An equal temperament is a musical temperament or tuning system, which approximates just intervals by dividing an octave (or other interval) into equal steps. This means the ratio of the frequencies of any adjacent pair of notes is the same, which gives an equal perceived step size as pitch is perceived roughly as the logarithm of frequency.

In classical music and Western music in general, the most common tuning system since the 18th century has been twelve-tone equal temperament (also known as 12 equal temperament, 12-TET or 12-ET; informally abbreviated to twelve equal), which divides the octave into 12 parts, all of which are equal on a logarithmic scale, with a ratio equal to the 12th root of 2 (12√2 ≈ 1.05946). That resulting smallest interval, 1⁄12 the width of an octave, is called a semitone or half step. In Western countries the term equal temperament, without qualification, generally means 12-TET.

In modern times, 12-TET is usually tuned relative to a standard pitch of 440 Hz, called A440, meaning one note, A, is tuned to 440 hertz and all other notes are defined as some multiple of semitones apart from it, either higher or lower in frequency. The standard pitch has not always been 440 Hz. It has varied and generally risen over the past few hundred years.(4,5)
